# Acoustic context modulates natural sound discrimination in auditory cortex through frequency specific adaptation

**DOI:** 10.1101/2021.02.08.430293

**Authors:** Luciana López-Jury, Francisco García-Rosales, Eugenia González-Palomares, Manfred Kössl, Julio C. Hechavarria

**Affiliations:** Institute for Cell Biology and Neuroscience, Goethe University, 60438, Frankfurt am Main, Germany

## Abstract

Vocal communication is essential to coordinate social interactions in mammals and it requires a fine discrimination of communication sounds. Auditory neurons can exhibit selectivity for specific calls, but how it is affected by preceding sounds is still debated. We tackled this using ethologically relevant vocalizations in a highly vocal mammalian species: Seba’s short-tailed bat. We show that cortical neurons present several degrees of selectivity for echolocation and distress calls. Embedding vocalizations within natural acoustic streams leads to stimulus-specific suppression of neuronal responses that changes sound selectivity in disparate manners: increases in neurons with poor discriminability in silence and decreases in neurons selective in silent settings. A computational model indicates that the observed effects arise from two forms of adaptation: presynaptic frequency specific adaptation acting in cortical inputs and stimulus unspecific postsynaptic adaptation. These results shed light into how acoustic context modulates natural sound discriminability in the mammalian cortex.

## Introduction

Studying how vocalizations are processed in the brain of listeners is key to understand the neurobiology of acoustic communication. How neurons respond to natural sounds has been studied in many animal species from primates to bats (Newman & Wollberg, 1973; Suga *et al*., 1978; Feng *et al*., 1990; Doupe & Konishi, 1991; Rauschecker *et al*., 1995; Esser *et al*., 1997). It is known that-at least some-neurons in the auditory system respond more to specific sound types. This so-called “natural sound selectivity” appears to be linked to temporal and spectral cues in the sounds perceived (Lewicki & Konishi, 1995; Wang *et al*., 1995; Schnupp *et al*., 2006; Liu & Schreiner, 2007; Ter-Mikaelian *et al*., 2013). Though natural sound selectivity has been studied extensively, we still debate whether and how it is affected when hearing in natural environments in which individual sounds are often preceded by other acoustic elements.

Sensory history refers to stimuli (auditory or from other modalities) occurring before a specific event. In the auditory modality, numerous studies have shown that leading sounds can modulate responses to upcoming ones. This modulation is present in phenomena such as forward masking (Brosch & Scheich, 2008; Scholes *et al*., 2011), stimulus specific adaptation (Nelken & Ulanovsky, 2007; Hershenhoren *et al*., 2014; Malmierca *et al*., 2014) and combination sensitivity (Margoliash & Fortune, 1992; Fitzpatrick *et al*., 1993; Esser *et al*., 1997; Macias *et al*., 2016), to name a few. One common issue found in the existing literature on auditory processing is that studies on neural selectivity to natural sounds often do not assess the influence of acoustic history on the selectivity observed (Klug *et al*., 2002; Medvedev & Kanwal, 2004; Grimsley *et al*., 2012; Mayko *et al*., 2012). Likewise, studies on how acoustic context modulates responses to upcoming sound events are, for the most part, conducted using artificial stimuli that often do not carry any ethologically relevant information for the animals (Ulanovsky *et al*., 2003; Scholl *et al*., 2008; Antunes *et al*., 2010; Phillips *et al*., 2017). One unfortunate result of this discrepancy is that, at present, we do not know how theories formulated in experiments that used artificial sounds apply when natural vocalizations are being listened.

To study the latter, we conducted a series of experiments to assess natural sound processing in the auditory cortex (AC) of awake bats (*Carollia perspicillata*). Specifically, we probed cortical responses to two very distinct sound types: echolocation (used for navigation in bats) and communication (represented here by distress sounds). While listening to echolocation signals indicates the presence of a navigating conspecific, hearing a distress call indicates the presence of an individual under duress and thus a potentially harmful situation. In bats, distress sounds have strong communicative power and they trigger autonomic (Hechavarria *et al*., 2020) and hormonal responses in the listeners (Mariappan *et al*., 2016). Navigation and distress sounds differ markedly not only in their ethological value but also in their acoustic attributes, especially in their frequency composition with distress energy peaking at ~23 kHz and echolocation peaking at frequencies above 60 kHz (Hechavarria *et al*., 2016).

Even though bat navigation and distress vocalizations are clearly different in their acoustics, there are neurons in the auditory system of *C. perspicillata* that can potentially respond to both signal types (Kossl *et al*., 2014; Gonzalez-Palomares *et al*., 2021). In other bat species, cortical neurons are known to respond to both navigation and social calls (Ohlemiller *et al*., 1996; Esser *et al*., 1997; Kanwal, 1999). These are neurons with ‘multipeaked’ or broad frequency receptive fields, i.e. when tested with pure tones they respond to both low (~ 20 kHz) and high (> 60 kHz) frequency tones. Studies in several animal models have shown that the peaks in ‘multipeaked’ frequency tuning curves often correlate with the spectral components of species-specific vocalizations (Kadia & Wang, 2003; Moerel *et al*., 2013; Kikuchi *et al*., 2014). Although the general consensus is that ‘multipeaked’ neurons are potentially useful for processing acoustic signals with complex spectral structures (Sutter & Schreiner, 1991), their broad tuning could make these neurons worse in discriminating between vocalization types that are potentially linked to different behaviours. To test this hypothesis, in the first part of this study we recorded activity of single AC neurons in response to natural distress and echolocation sounds. We show that-as hypothesized-the majority of multipeaked neurons can discriminate poorly between distress and echolocation signals. However, there exist as well neurons that respond equally well to distress and navigation calls albeit having single-peaked frequency tuning curves.

In the second part of the study we tested how acoustic context (i.e. previous history) affects cortical responses to echolocation and distress sounds. To that end, navigation and distress sounds were presented preceded either by sequences of other navigation or distress vocalizations, resulting in stimulation conditions with expected and unexpected acoustic transitions, e.g. a distress sound following navigation calls would be “unexpected”. This experimental design aimed to mimic complex acoustic transitions that animals encounter in their natural environment. Our hypothesis was that leading acoustic context could modulate responses to upcoming sounds. We predicted responses to expected sounds to be diminished and responses to unexpected sounds to be either enhanced or less suppressed. This idea is based on the rich body-of-knowledge indicating that AC neurons are context sensitive as proven using, among others, stimulus specific adaptation (and forward masking paradigms (Brosch & Scheich, 2008; Scholes *et al*., 2011; Hershenhoren *et al*., 2014; Malmierca *et al*., 2014). The results obtained confirmed our hypothesis: leading acoustic context drives stimulus-specific suppression on the cortical responses to natural sounds with the strongest suppression occurring in response to expected sounds. To our surprise, adding acoustic context before the target sounds turned poor discriminator neurons into good distress-vs.-navigation discriminators, and good discriminators into poor ones. In other words, when studied in acoustically-rich and ethologically-relevant situations, natural sound discrimination at the single neuron level behaves radically different from experiments in which single sounds are used as stimuli.

To assess the possible origins of these results, in the third part of the study, we built a computational neuron model that was capable of replicating experimental observations. The model predicted the existence of two adaptation mechanisms: one operating at the presynaptic level that contributed to the context-specific suppression observed; and another one operating at the postsynaptic level (i.e. directly on cortical neurons) that supressed spiking output in a context-unspecific manner. This model shares similarities with frequency-specific adaptation models used to explain stimulus-specific adaptation and the processing of speech sounds in humans (Mill *et al*., 2011; Taaseh *et al*., 2011; May *et al*., 2015). The results of the present study shed light onto the neural basis of natural sound processing in the auditory cortex, and link natural sound discrimination to acoustic-context effects on sensory processing in a highly vocal mammal.

## Results

We investigated how auditory cortex neurons of the bat *C. perspicillata* process two types of natural vocalizations: echolocation and communication calls. The goal was to assess how leading sequences of behaviourally relevant categories, echolocation and communication vocalizations, affect cortical responses to lagging sounds (called probes throughout the text, see Fig. 1). Probe sounds could be either *expected* (e.g. an echolocation syllable occurring after an echolocation sequence, Fig. 1c) or *unexpected* (e.g. distress syllable following an echolocation sequence). Note that this terminology refers only to neural expectations and not to a cognitive process. Probes were presented 60 ms after the context offset. In *C. perspicillata*, the main energy of echolocation and communication calls occurs at different frequencies: ~66 kHz in echolocation calls (Fig. 1b, second column) and ~23 kHz in distress calls (Fig. 1b, fourth column). In addition to their frequency composition, echolocation and distress sequences differ in their temporal structure (Fig. 1b first and third columns).

**Figure 1.**
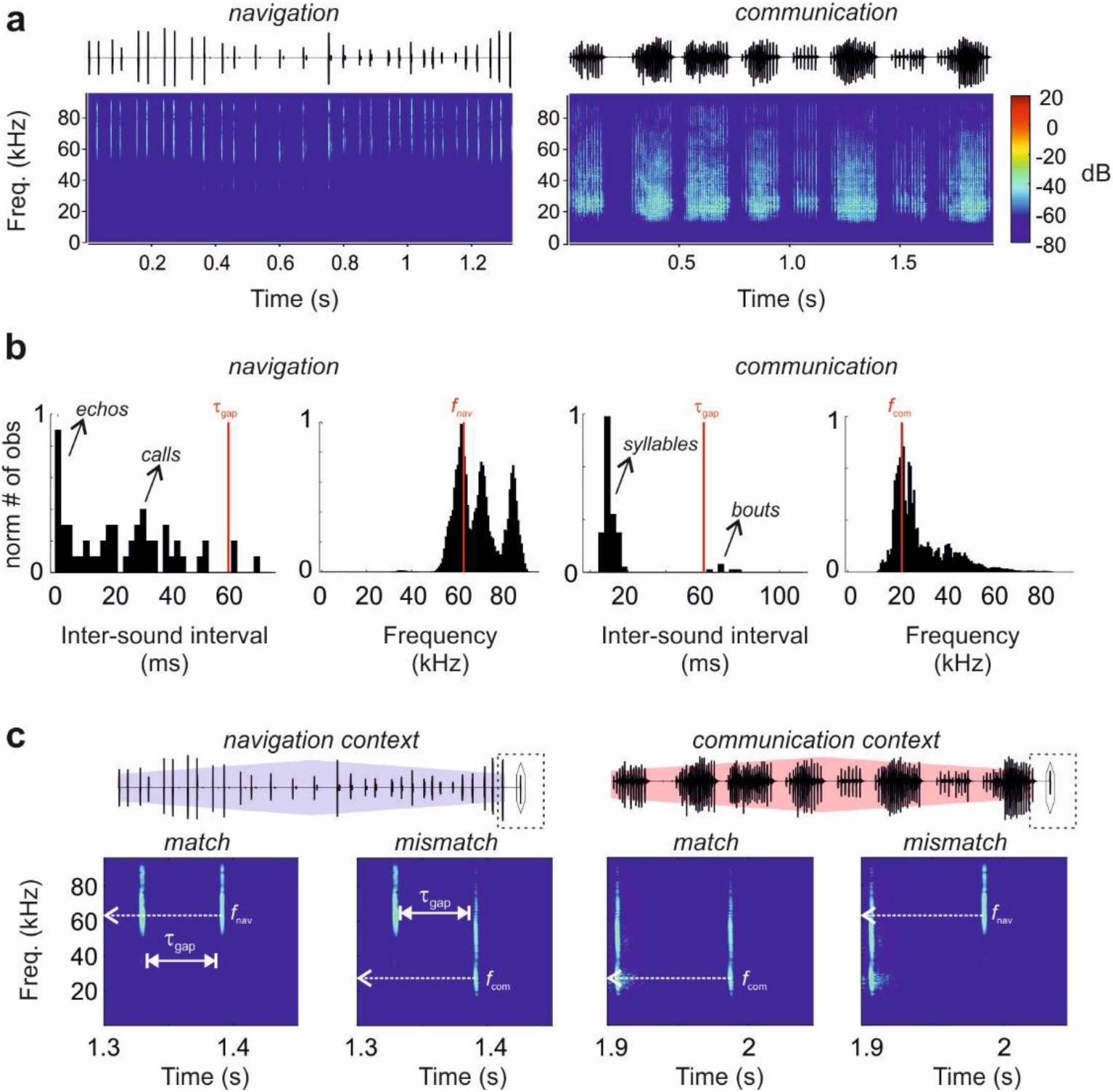
The stimulation paradigm. **a** Oscillogram and spectrogram of natural calls of *C. perspicillata* associated with navigation and communication behaviours: a sequence of echolocation calls (left) and a distress call (right). **b** Distribution of inter-sound intervals and frequencies of the sequences above. Arrows indicate independent components within the sequences. The vertical red lines in temporal distributions indicate the temporal gap between the end of the natural sequence and the onset of the probe in our stimulation paradigm (τ_gap_ = 60 ms). In spectral distributions, red lines indicate the peak of the distribution (*f*_nav_ = 66 kHz and *f*_com_ = 23 kHz). **c** Oscillograms of the stimulation protocol. Natural sequences preceding a short probe (highlighted by a diamond). Below, spectrograms of the last 150 ms of stimulus (dashed rectangle) point out the last element of each sequence and the subsequent probe. Time between the offset of the sequence and the onset of the probe is indicated as τ_gap_ in white and the frequency at peak energy of the probes is indicated by dashed arrows.

### Cortical responses to echolocation and *communication* syllables after silence

The extracellular activity of 74 units was recorded in the auditory cortex (AC) of awake bats. The neurons were classified into five groups regarding their responses to the probes presented in isolation, i.e. preceded by 3.5 s of silence (see Methods for details). The majority of units (91%, 67 out of 74) responded to both sounds (grey bars in Fig. 2a), the remaining 9% (7/74 units) responded exclusively to echolocation or communication (black bars in Fig. 2a). Note that we selectively targeted high frequency regions of the auditory cortex that respond to both low and high frequencies (Hagemann *et al*., 2010; Hagemann *et al*., 2011) and thus this result was expected.

**Figure 2.**
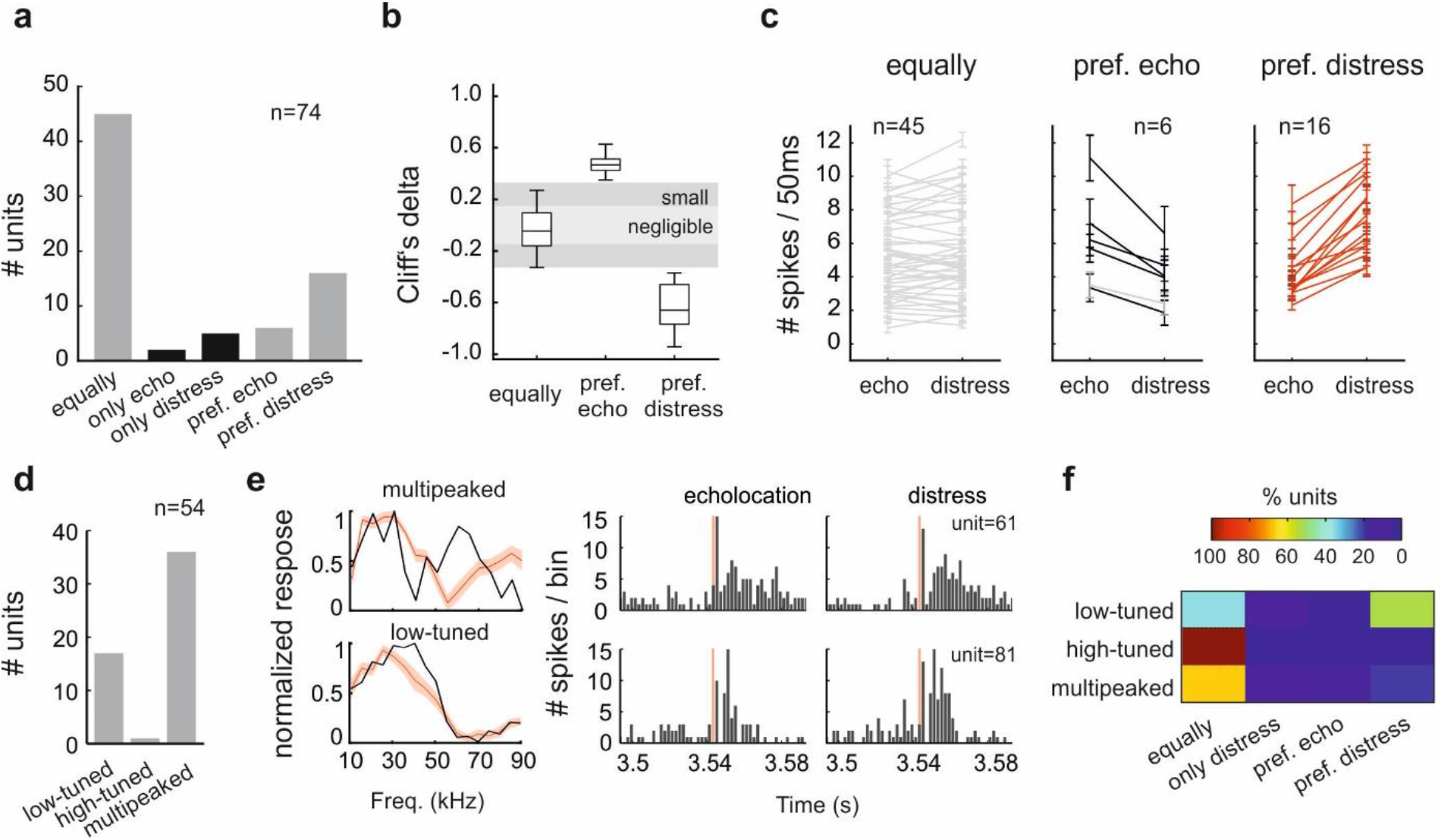
Neuronal classification of the responses to natural sounds without acoustic context. **a** Number of units classified according to their responses to a single echolocation call and a distress syllable. Categories were defined considering the number of spikes evoked during 50 ms from the sound onset (‘equally’: abs(Cliff’s delta)<=0.3; ‘pref.’: abs(Cliff’s delta)>0.3). Grey bars correspond to units responsive to both sounds. Black bars, to units that are only responsive to one sound. **b** Cliff’s delta calculated for neuronal classification for units responsive to both sounds. Size effect interpretation are indicated by shade areas. **c** Average number of spikes across trials in response to both probes per classification. Grey lines indicate non significative differences between spike counts (*p*-value>0.05, Wilcoxon rank sum test). Red and black lines correspond to units that showed significative differences (*p*-value<0.05, Wilcoxon rank sum test). Error bars correspond to standard errors across trials. **d** Number of units according to their iso-level frequency tuning shape. Low- and high-tuned correspond to one-peak tuning curves, peaking at <50 kHz and >50 kHz, respectively. **e** Two examples of units classified as ‘multipeaked’ (black top) and low-frequency tuned (black bottom). The red iso-level tuning curves indicate the average across units and the shade area, the standard error. At the right, the corresponding peri-stimulus time histogram (PSTH) in response to each probe. **F**, Percentage of units classified by their response to natural sounds per iso-level tuning curve shape.

In each unit, the number of spikes evoked by the two probes-that is, single echolocation and communication syllables-were statistically compared using the Cliff’s delta metric. Units with *negligible* and *small* effect size (abs(Cliff’s delta)<=0.3) were defined as ‘equally’ responsive to both sounds (45/74 units). Otherwise, units with larger effect size were classified as ‘preference for communication (16/74 units) or ‘preference for echolocation’ (6/74 units) by comparing their mean spike counts (Fig. 2b). Fig. 2c shows the spike count of all units split into the three groups mentioned.

### Linking frequency receptive fields to natural sound responsivity

Next, we tested whether the units’ responsiveness could be explained on the basis of overlap between the frequency spectrum of call components and their iso-level frequency tuning curves measured at 60 dB SPL. Iso-level frequency tuning curves were calculated using pure tones covering frequencies from 10 to 90 kHz (5 kHz steps) for a subset of neurons (n = 54 out of 74 units). Their classification according to the shape of the iso-level tuning curve is shown in Fig. 2d (see Methods for the procedure). Sixty-seven percent of the units (36/54 units) presented multipeaked frequency tuning curves (example unit in top Fig. 2e). This feature has been associated to non-tonotopically arranged areas in the AC (high-frequency fields and dorsoposterior area) in *C. perspicillata* (Hagemann *et al*., 2010; Lopez-Jury *et al*., 2020). The remaining units studied (18/54) exhibited single-peaked tuning, associated to neurons in tonotopic areas (primary and secondary AC and anterior auditory field). From those, 17 were low-frequency tuned (best frequency < 50 kHz, example unit in bottom Fig. 2e) and only one had best frequency >= 50 kHz. The iso-level tuning curves of all multipeaked and low-frequency tuned units are shown in Supplementary Fig. 1 and the respective mean curves in Fig. 2e.

As expected, the majority of the neurons with multipeaked tuning curves (67%) belonged to the group ‘equally responsive’ (example in Fig. 2e, top). In addition, 65% of the neurons with low-frequency iso-level tuning curve presented preferential or exclusive responses for communication sounds (example in Fig. 2e, bottom). These results indicate that, though simple, iso-level tuning curves partially predict neuronal responsiveness to natural sounds. The percentages of responsivity to natural sounds for each tuning curve group are shown in Fig. 2f.

### Natural acoustic context supresses cortical responses to probe sounds

The effect of acoustic context-i.e. navigation or communication sound sequences preceding the presentation of probe sounds-was quantified for the 45 units that responded equally well to both probes when presented after silence (Fig. 2c, left). Fig. 3a depicts the PSTHs of an example unit in response to the echolocation and communication syllables preceded either by silence (top), by a navigation context (middle) or by a communication context (bottom). In this unit, the response to both probes was reduced after context. In fact, the response to the echolocation call was completely abolished after the navigation context (Fig. 3a, middle panel left). Overall, the presence of a leading context decreased responsivity in the majority of cortical neurons regardless of the type of probe sound that followed the masking sequences (N values between 36-29 out of 45 units, *p* < 0.05, rank sum). Note that all black lines (significant changes) point downwards in Fig. 3b1. However, a subset of neurons did not present significant variations in the response to the probe after context (*p* > 0.05, rank sum), the respective spike-counts are plotted with grey lines in Fig. 3b1 (N values between 9-16).

**Figure 3.**
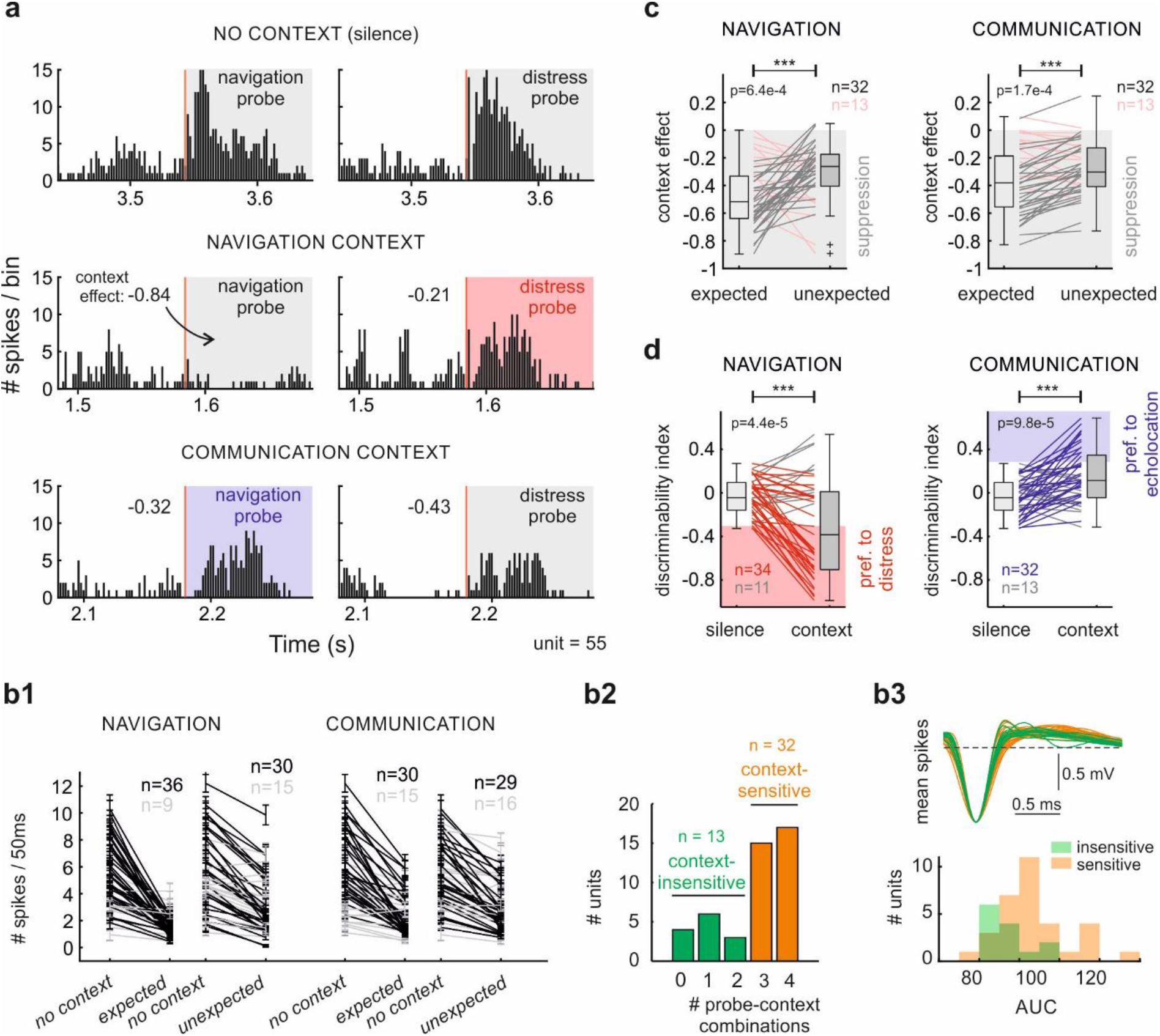
Stimulus-specific suppression on neuronal responses by leading acoustic context. **a** Responses to echolocation call and distress syllable of a unit classified as ‘equally responsive’, after 3.5 s of silence (top), after a sequence of echolocation (middle) and after a sequence of distress (bottom). **b1** Average number of spikes across trials in response to a probe after silence (*no context*) and after context (*expected* or *unexpected*). *Expected* in a navigation context corresponds to the response to an echolocation call and *unexpected* to a distress syllable; and vice-versa in the communication context. Black lines correspond to significant decrement (*p*-value<0.05, Wilcoxon signed rank test) and grey, non-significant. Error bar corresponds to standard error across trials. **b2** Number of units that presented significative suppression in 0 to 4 context-probe combinations. **b3** Spike waveforms (average per isolated single unit) of context-insensitive neurons (green) and context-sensitive neurons (orange). At the bottom, the distributions of the corresponding areas under the waveform. **c** Context effect quantification of ‘equally responsive’ units for *expected* and *unexpected* probes. Lines join values from the same neuron. Grey lines correspond to units with a higher suppression on *expected* sounds. Otherwise, the lines are pink. **d** Cliff’s delta of ‘equally responsive’ units comparing echolocation and distress responses after silence and after acoustic context. Significance level is indicated above the plots c-d; *p*-values were obtained using the paired test Wilcoxon signed rank.

Thirty-two units showed significant reduction of the response in at least three context-probe combinations and were considered as ‘context-sensitive’ neurons. The rest of the neurons (n=13) were classified as context-insensitive (Fig. 3b2). In order to study if these two classes of neurons differed from each other on their intrinsic properties, we compared best frequency, spontaneous firing rate and spike waveform. Only the shape of the spikes was significantly different between the two groups (Fig. 3b3). Context-sensitive neurons showed higher values of area under the spike waveform than context-insensitive, suggesting that context-sensitive neurons could be classified as broad-spiking putative pyramidal cells and the context-insensitive neurons, as narrow-spiking putative interneurons (Tsunada *et al*., 2012).

### Context increases cortical discrimination of natural sounds

Although the example in Fig. 3a shows reduction of the probe-responses by leading context, the magnitude of the suppression was different when comparing between probes. The *expected* probes (the ones matching the leading context) in both context categories were more suppressed than the *unexpected* probes relative to the responses after silence (see context effect suppression values on top of each PSTH). The context-dependent effect illustrated in Fig. 3a was not unique to this example neuron. In fact, stronger reduction of responses to *expected* probes were common among the neurons studied (Fig. 3c), regardless of the type of context that preceded the probes (*p* = 6.4e-4 for navigation and *p* = 1.7e-4 for communication, signed rank). Relative to the probes preceded by silence, there was a median of 48 % of suppression for *expected* sounds versus 30 % for *unexpected* after navigation context, and 37% and 28% for communication context, respectively.

So far, we have shown that non-selective neurons for natural sounds of different behavioural category-navigation and communication-exhibit suppression by leading acoustic context that is stimulus-specific. We reasoned that such context-specific suppression could actually render cortical neurons more “selective” to natural sounds than what one would predict by presenting single sounds preceded by silence. To quantify if this was the case, we compared the Cliff’s delta values (used here as a discriminability index) obtained when comparing spike counts between the isolated probes, versus the value obtained when there was leading context (Fig. 3d). After both contexts, the discriminability of the probes significantly increased (values closer to −1 or 1) compared to the discriminability exhibited in silence (*p* = 4.4e-5 after navigation and *p* = 9.8e-5 after communication, signed rank). The results showed that under navigation context, Cliff’s delta values became more negative, from a median of −0.045 (in silence) to – 0.38, indicative of higher number of spikes in response to communication syllable. On the other hand, the presence of communication context shifted Cliff’s delta values closer to 1 (from a median of −0.045 to 0.11), which corresponds to a higher number of spikes in response to echolocation call. Similar results were obtained when the delay between probe and context was set to 416 ms (Supplementary Fig. 2a).

Overall, these results show that, as hypothesized, the responses to communication or echolocation syllables presented alone are more difficult to discern than when the syllables are presented after a behaviourally relevant acoustic sequence. In other words, acoustic contexts decreased the ambiguity of the response of non-selective cortical neurons by increasing the sensitivity to temporal transitions between these two sound categories.

### Effects of acoustic context on neurons with preference for *communication* sounds

We also examined the effects of context sequences on the response of neurons classified as ‘preference for communication’ sounds. These neurons formed the second largest neuronal group observed when presenting the sounds without context (22% of the neurons studied, Fig. 2c, right). A typical example neuron that ‘preferred’ communication syllables can be observed in Fig. 4a (top). Note that although this neuron has a stronger response to the communication syllable, it still responded well to the echolocation call presented without any acoustic context. In this example, as in the previous neuronal group (‘equally responsive’ neurons, Fig. 3), the leading context suppressed the response to both probes and the suppression was stronger on *expected* sounds (second and third row in Fig. 4a). The responses to the probes after context were analysed for all the ‘preference for communication’ units (n=16) and significant suppression after context was found in the majority of the neurons (Fig. 4b). Moreover, stimulus-specific suppression was found also at the population level (Supplementary Fig. 3), following the same trend exhibited by the example unit in Fig. 4, as well as in the ‘equally responsive’ neurons group (Fig. 3c).

**Figure 4.**
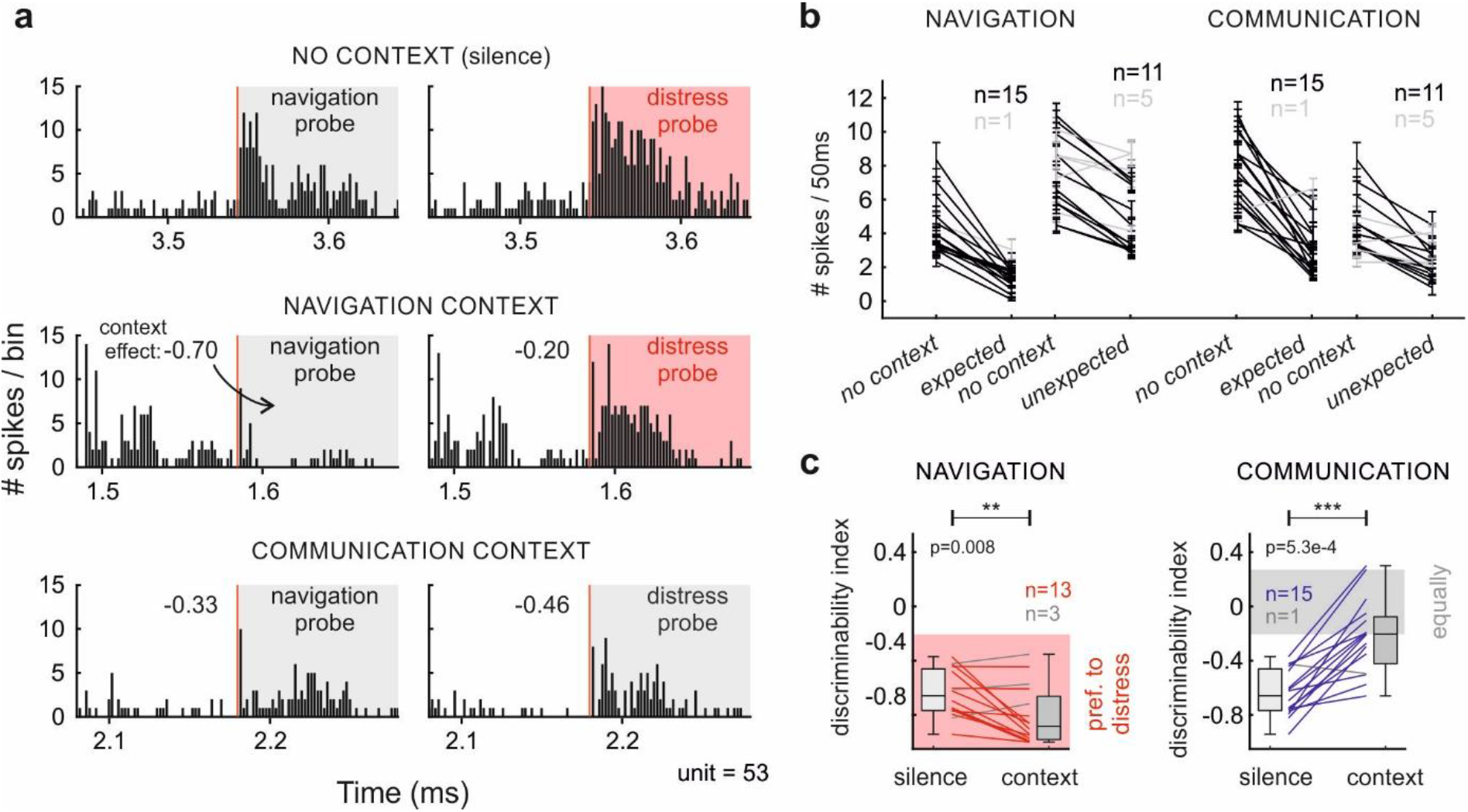
Context modulation in selective neurons decreased neuronal discriminability. **a** Responses to echolocation call and distress syllable of a unit classified as ‘preference for distress’, after 3.5 s of silence (top), after a sequence of echolocation (middle) and after a sequence of distress (bottom). **b** Average number of spikes across trials in response to a probe after silence (*no context)* and after context (*expected* or *unexpected*). Black lines correspond to significant decrement (*p*-value<0.05, Wilcoxon rank sum test) and grey, nonsignificant. Error bar corresponds to standard error. **c** Cliff’s delta of ‘preference for distress’ units comparing responses to echolocation and distress syllables after silence and after acoustic context (Wilcoxon signed rank test).

The clearest difference between these selective neurons versus the non-selective ones occurred when the probe sounds were preceded by the communication context. To illustrate, in the example in Fig. 4a (bottom), when preceded by a communication context, the responses to communication and echolocation syllables became similar, even though they were notably different when the sounds were presented without context (Fig. 4a, top). This result stems from the unbalanced suppression on probe-evoked responses (−0.46 on *expected* versus −0.33 on *unexpected*) together with the intrinsic neuronal preference for communication sounds, that brought spike outputs in response to each probe to the same level when they were presented after the communication sequence.

Equally strong responses to the probes after communication context was also present at the population level. To quantify the difference between probe-evoked responses, we calculated Cliff’s Delta values of the spike counts in response to both probes after silence and sequences (Fig. 4c). In agreement with the example, when the context was communication, Cliff’s delta values went from very negative values (preference for communication when preceded by silence) to higher values (*p* = 5.3e-4, signed rank), closer to zero, i.e. similar probe-evoked responses, (Fig. 4c right, see Supplementary Fig. 2b for data obtained with 416-ms delays). In addition, when the context was the navigation sequence, Cliff’s Delta values became even more negative after context (*p* = 0.008, signed rank), indicating better discrimination and higher responsivity to *unexpected* (communication) sounds. These results indicate that good neuronal navigation-communication classifiers-according to spike counts measured with single sounds presented in silence-became in fact poor category discriminators when tested in a context of communication sounds.

### A computational model with frequency specific adaptation explains context dependent suppression observed in-vivo

While single navigation and communication syllables evoked similar responses in most cortical neurons, after acoustic context, the responses shifted towards the syllable that corresponded to the *unexpected* sound. To identify which mechanisms can account for these context-dependent effects, we implemented a model of an ‘equally responsive’ cortical neuron based on leaky integrate-and-fire neuron models.

First, we modelled a single cortical neuron, set to be ‘equally responsive’ to both sounds categories when presented alone, and with two synaptic inputs whose firing are highly selective for either communication or echolocation signals (see diagram in Fig. 5a1). Each synapse receives spike trains whose firing rate is proportional to the degree of selectivity that each input was assumed to have for each sound category. The temporal pattern of the spike trains was assumed to follow the envelope of the respective natural sound used in each simulation. We also assumed that the degree of the input selectivity is correlated with the spectral components of the natural sounds used in our experiments. Thus, we distinguished two input classes: high-frequency tuned (> 45 kHz) and low-frequency tuned (< 45 kHz). The rate of the inputs in response to each syllable are shown in Fig. 5a1. Note that the firing rate of both inputs increases in response to the communication syllable, albeit to a higher degree in the low-frequency tuned input. The latter is a consequence of the spectral broadness of communication calls that carry energy in both low and high frequency portions of the spectrum, as opposed to navigation signals that, in this bat species, are limited to frequencies above 50 kHz.

**Figure 5.**
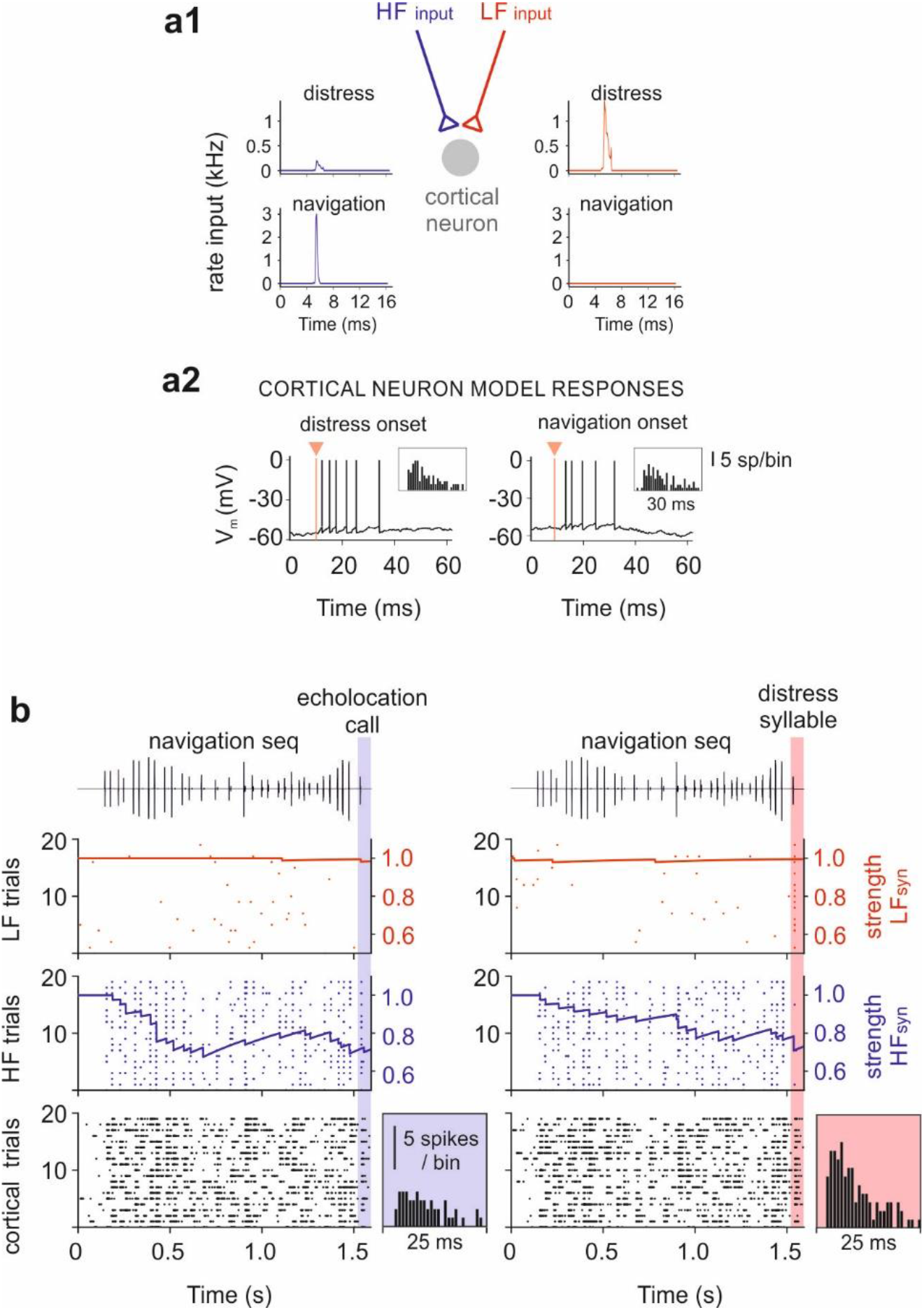
Computational modelling of stimulus-specific suppression by leading context on cortical neurons. **a1** Diagram of the components of the model. Red and blue arrows correspond to the inputs of the neuron model. The rate (in kHz) of the spiking of each input in response to echolocation call and distress syllable. **a2** Membrane potential of the neuron model in response to both probes. Insets show simulated PSTH across 20 trials for the corresponding sounds. **b** Example of 20 simulations of neuronal responses to a navigation sequence followed by an echolocation call (left) and by a distress syllable (right). Raster plots indicate the spiking of the inputs (top and middle) and of the cortical neuron model (bottom). On top of the rasters, the time evolution of the synaptic strength associated to the spiking of the first trial of each input. The insets show the PSTH obtained from the neuron model during 25 ms after the onset of the probe for each stimulation. The parameters used in the model are indicated in Table 1.

In the neuron model, each presynaptic spike induces an increment of excitatory conductance. The amplitude of this conductance change was adjusted so that the neuron responded equally to both probe sounds (echolocation and communication sounds preceded by silence), as observed in the majority of neurons measured in this study. Equivalent spiking responses to both probe sounds in one trial simulation are illustrated in Fig. 5a2. We ran 20 trial simulations per probe stimulation and the resultant PSTHs are shown in the respective insets of Fig. 5a2.

Simulated responses to the navigation context followed by an echolocation call and by a communication syllable are shown in Fig. 5b. The spiking of the low-frequency tuned input (red), high-frequency tuned input (blue), and that of the cortical neuron (black) are shown as raster plots in the top, middle and lower panels, respectively. Looking at the responses to the navigation context in this simulation (i.e. times before the 1.5-s mark), we observe that while the spiking of the low-frequency input (top) across 20 trials corresponds only to spontaneous activity of the input, the spiking of the high-frequency channel (middle) tightly follows the envelope of the echolocation sequence. As mentioned in the preceding text, this input selectivity was built in the model, and it is only described here to illustrate how the model works. The spiking of the cortical neuron (bottom) also exhibited a response to the navigation sequence. The latter is also expected since cortical neurons integrate both low and high frequency inputs.

To replicate *in-vivo* results *in-silico*, we implemented activity-dependent adaptation in the model. A likely mechanism of adaptation is synaptic depression and it has been proposed to underlie context-dependent processing (Ulanovsky *et al*., 2004; Wehr & Zador, 2005). The temporal course of the synaptic strength associated to the spiking of the first simulation trial of each input is depicted in solid lines on top of the corresponding raster plots (Fig. 5b). The synaptic strength is proportional to the amplitude of the excitatory postsynaptic potential that each presynaptic spike produces in the cortical neuron; low values indicate less probability of spiking, high values the opposite. Because synaptic depression is activity-dependent, the synapse that receives activity from the high-frequency tuned input is more adapted (lower values of synaptic strength) than the synapse that receives only spontaneous activity from the low-frequency tuned input. The recovery time of the adaptation implemented allowed to affect output responses to forthcoming probe sounds 60 ms after the offset of the context sequence. Consequently, the spiking associated to the echolocation probe in the high-frequency tuned input (blue area in the middle-left raster plot) generated less spikes in the cortical neuron than the spiking in the low-frequency tuned input in response to the communication syllable (red area in the top-right raster plot). The latter is explained by lower values of synaptic strength associated to the high-frequency tuned synapse in comparison with the low-frequency tuned synapse at the probe onset. To illustrate the difference between cortical output in response to both probes after a navigation sequence, we plotted the respective PSTHs on the right of each raster plot (Fig. 5b bottom). Note that in the absence of context, this neuron model responded equally strong to both probe sounds because both synapses are equally adapted (see Fig. 5a2). A previous and constant stimulation (echolocation sequence) in the model unbalances the synaptic adaptation, prevents equal responses to communication and echolocation probes, and replicates the results obtained *in-vivo*.

Simulations performed with the “communication context” can be found in Supplementary Fig. 4. Note that in this case, we were able to replicate *in-vivo* results only after decreasing the magnitude and the recovery time of the synaptic depression associated to the spiking of low-frequency tuned input (see Methods section and Supplementary Fig. 5 for details).

### Requirements of the model to reproduce data

Our model suggests that stimulus-specific context effect arises out of (1) segregated inputs selective for each sound category and (2) synaptic adaptation. To illustrate that these conditions are necessary to reproduce our empirical results, we ran different simulations modifying the parameters associated to these properties.

First, we varied the degree of input selectivity and compared the Cliff’s delta between spike counts obtained in response to navigation versus communication probes (discriminability index), as in our data analysis (Fig. 6a). The simulations showed that when the inputs have no preference for any of the signals (both inputs respond to both), the discriminability index did not change significantly after context (*p* > 0.05, rank sum; grey boxplots in Fig. 6a). These results proved that input selectivity was necessary to obtain stimulus-specific context effects. On the other hand, if the inputs were selective exclusively to one sound category (either navigation or communication), the discriminability index was significant and drastically different after context compared with silence (*p* < 0.001, rank sum; green boxplots in Fig. 6a), following the trend found empirically. However, only simulations with navigation context showed no statistical differences with our data (*p* = 0.3 navigation; *p* = 6.0e-11 communication, rank sum). A more biologically plausible model would assume inputs responding to the spectral features of sounds and not to sound categories: probably a middle point between no-selectivity and high selectivity. Indeed, the latter model rendered significant changes after context (*p* < 0.001, rank sum; orange boxplots in Fig. 6a), but a less dramatic effect than in the previous model with high selectivity. That is, when cortical inputs showed intermediate level of selectivity, simulations with both contexts showed no differences with the empirical data (*p* = 0.1 navigation; *p* = 0.7 communication, rank sum). Note that in this case, the input tuned to high frequencies would respond at least to some extent to the high frequency components found in communication signals.

**Figure 6.**
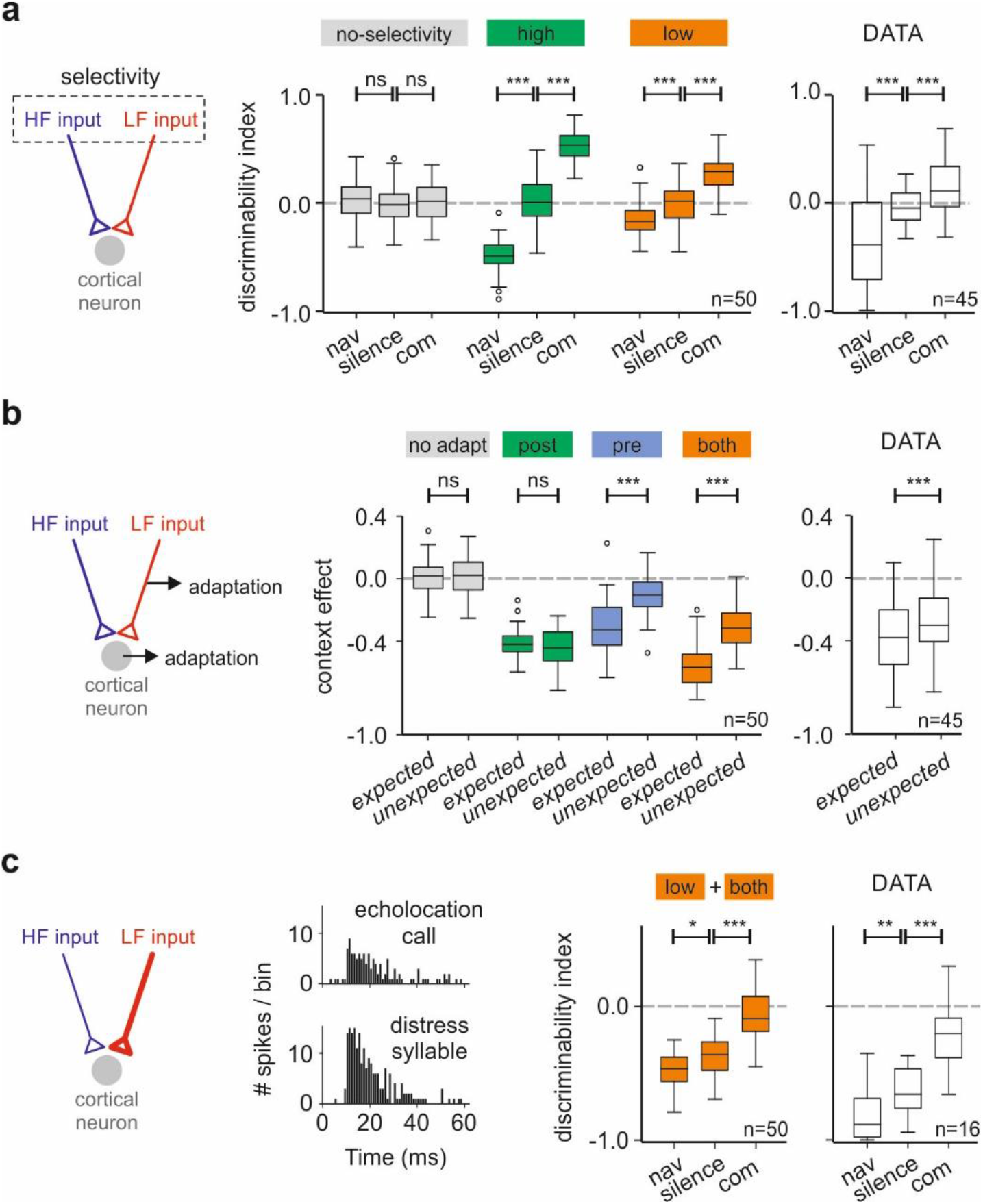
Requirements of the model to reproduce empirical results. **a** Discriminability index after silence, after the echolocation sequence (nav) and after distress (com) using models with different degrees of selectivity in the inputs. ‘No-selectivity’: model in which inputs were not selective for any type of signals, navigation or communication, although responded to both of them. ‘High’: inputs exhibited exclusive selectivity for one signal. ‘Low’: input selectivity correlates with the spectral components of the natural sounds. Right boxplots show the discriminability index obtained from our empirical data. **b** Communication context effect calculated for *expected* and *unexpected* probes using models with different forms of adaptation. ‘No adapt’: model without any type of adaptation. ‘Post’: model with neuronal adaptation. ‘Pre’: model with synaptic adaptation. ‘Both’: model with neuronal and synaptic adaptation. Right boxplots show the respective context effect obtained from experimental data. **c** Simulated PSTHs in response to both probes and discriminability index obtained from a model in which the synaptic weights were unbalance (as illustrated in the diagram). The neuron model assumes ‘low’ and ‘both’ conditions. In all the comparisons, the significance level is indicated above; *p*-values were obtained using non-paired Wilcoxon rank sum test for the simulations, and the paired Wilcoxon signed rank for the real data. The parameters used on each model are indicated in Table 2.

Secondly, we tested how different forms of adaptation influence the context-dependent modulation observed in our data. Fig. 6b shows the results of simulations using the communication sequence as leading context. As expected, a model that lacks any type of adaptation showed that the context effect on both probes was null (grey boxplots in Fig. 6b). Otherwise, a model that includes neuronal adaptation, dependent on postsynaptic spiking activity, displayed suppression of around 45% on both probes that were not significantly different from each other (green boxplots in Fig. 6b). This indicates that adaptation at the cortical neuron level allows context suppression, but not stimulus specificity. Alternatively, if the adaptation depends on presynaptic activity, the context effect becomes stimulus-specific (significant differences between the probes in blue boxplots in Fig. 6b). However, the response to the *unexpected* sound after navigation context is not suppressed (*p* > 0.05, one-sample rank sum, see Supplementary Fig. 6), which disagrees with our experimental data (Fig. 3c). On the other hand, a model that includes adaptation both at the level of the cortical neuron and at the presynaptic level rendered a suppressed response to *unexpected* sounds after navigation context (*p* < 0.05, one-sample rank sum) reproducing all the effects observed *in-vivo*. In other words, synaptic adaptation is necessary to create stimulus-specific suppression, but not enough to explain a reduction of the responses to *unexpected* sounds. We used the same models to run simulations with the echolocation sequence set as context and found similar results (see Supplementary Fig. 6).

### Modelling responses in neurons that preferred communication sounds

So far, we have shown that our computational model successfully explains the increment of neuronal discriminability after acoustic context observed in the majority of the neurons, i.e. those that were equally responsive to both echolocation and communication sounds when preceded by silence. However, a smaller set of neurons whose response to communication sounds was stronger than to echolocation, showed a decrease in the discriminability index after the communication context. To determine whether the same neuron model could explain the behaviour of this subset of neurons, we modified the maximum synaptic weight (*w_e_*) of both synapses in order to obtain a neuron model that possesses selectivity for communication sounds. After increasing the *w_e_* of the low-frequency tuned synapse and decreasing the *w_e_* of the high-frequency tuned synapse, our neuron model presented stronger responses to communication stimulation than to echolocation (see PSTHs in Fig. 6c). Without changing any other parameters in the model, we calculated the discriminability index between probes after silence and after context. The results are shown in the left boxplots in Fig. 6c. Comparable with our data (right boxplots in Fig. 6c), the neuron model increased its discriminability for the probes after the navigation context (more negative Cliff’s delta values in comparison with silence), but decreased the discriminability after the communication context (Cliff’s delta values close to zero). To further illustrate this, we plotted the PSTHs in response to the probes across 20 simulation trials obtained after each context (Supplementary Fig. 7b). The results suggest that although these two classes of neurons (‘equally responsive’ and ‘preference for communication’) possess different degrees of selectivity for single isolated natural sounds, the effects of leading context on their responses to forthcoming sounds can be explained by the same mechanisms.

## Discussion

It is known that the auditory system is selective to natural and behaviourally relevant sounds. However, it is still under debate how the system detects and discriminate between specific calls among the large repertoire of animal vocalizations, especially when the calls occur within the context of other natural sounds. This study demonstrates that acoustic context affects neuronal discriminability to natural sounds in the auditory cortex of awake bats. The presence of acoustic context before the calls in question created disparate effects on neuronal sound discriminability. Context increases discriminability in units that are non-selective when tested in a silent condition (i.e. with no leading context) and has the opposite effect on units that are selective in the absence of context. An *in-silico* evaluation of our results using computational modelling predicts that two forms of adaptation, presynaptic- and postsynaptic-dependent, are necessary to reproduce the context-dependent effects *in-vivo*. Furthermore, we predict that the presynaptic activity of the context-sensitive neurons is tuned to the spectral components of the natural sounds. Our results shed light into the neuronal mechanisms of hearing in bats, but these mechanisms could be-by hypothesis-shared by other mammals.

### Cortical neurons process both echolocation and communication signals

Previous studies have hypothesized that neurons in HF fields of the bat auditory cortex process both sound types: navigation signals and communication calls (Ohlemiller *et al*., 1996; Esser *et al*., 1997; Kanwal, 1999; Kossl *et al*., 2014). Here, we provide strong evidence that corroborates this hypothesis. The majority of the neurons tested in this study (91%) presented responses to conspecifics’ echolocation calls and to a particular type of communication call, distress calls. In a previous study, we demonstrated that neurons in the inferior colliculus of the same bat, *C. perspicillata*, process the same two types of calls (Gonzalez-Palomares *et al*., 2021). In other mammalian species, cortical neurons can respond to multiple sound categories (Newman & Wollberg, 1973; Winter & Funkenstein, 1973; Tian *et al*., 2001; Grimsley *et al*., 2012), although the spectral differences between the sound types are not as extreme as those observed when comparing echolocation vs. communication in bats.

We show that more than two-thirds of the neurons that respond to both sounds categories are unable to discriminate the calls by means of rate coding. What this result means for natural hearing is that if a bat hears an isolated echolocation or communication call (i.e. with no context) most neurons in HF fields will fire and the spike output of most individual HF neurons will not allow the bat to discriminate the heard sound. This does not mean that the bat is not able to discriminate, as discrimination could still be achieved by means of temporal codes (Wang & Kadia, 2001; Schnupp *et al*., 2006; Liu & Schreiner, 2007; Huetz *et al*., 2011) or based on the activity of other AC neurons with less broad frequency tuning curves, such as those found in AI, AII or AAF. In the current study, we also report a fraction of HF fields neurons that were more responsive to one particular call (either echolocation or communication). Those neurons could also play an important role in the neural categorization of single isolated natural sounds.

A significant proportion of the neurons exhibited ‘multipeaked’ frequency tuning curves, which is consistent with the fact that we recorded mostly from HF fields (Hagemann *et al*., 2010; Hagemann *et al*., 2011). As expected, the majority of the ‘multipeaked’ neurons were equally responsive to both natural sounds used in our study. Additionally, the majority of low-frequency tuned neurons exhibited preference for sounds with higher power at low frequencies, the communication syllables. However, we found neurons (n = 6) that were barely responsive to pure tones at high frequencies (~ 60 kHz) and still presented responses to echolocation calls that were as strong as those obtained with communication calls. This discrepancy can be due to iso-level tuning curves not being able to predict responsivity to natural calls. Several studies have shown that the neuronal responses to complex sounds cannot be predicted based on its response profile to pure tones (Machens *et al*., 2004; Sadagopan & Wang, 2009; Laudanski *et al*., 2012; Feng & Wang, 2017). Bat neurons do not appear to be an exception.

### Acoustic context modulates natural sound discrimination in auditory cortex neurons

Our results provide evidence for a strong involvement of neurons in non-tonotopic areas in the processing of natural sounds (echolocation and communication) when these sounds are presented in isolation. Yet, bats are highly vocal animals and in a natural scenario, sonar and social calls from multiple conspecifics form an acoustic continuum in which sounds are often preceded by other sounds.

A well-established notion is that previous acoustic history (leading context) modulates neural activity in mammals (for review see Angeloni and Geffen (2018)). Generally, this is studied regarding the processing of artificial sounds, accounting for phenomena like SSA, forward masking and predictive coding (Calford & Semple, 1995; Auksztulewicz & Friston, 2016; Carbajal & Malmierca, 2018). Consistent with such studies, we observed that, overall, responses to *expected* sounds are more suppressed than responses to *unexpected* sounds.

This provides strong evidence indicating that the acoustic transitions between ethologically relevant sounds (communication and echolocation in bats) are represented in auditory cortex neurons in a manner similar to what has been postulated in studies that used artificial stimuli.

The data presented in this paper advances our thinking on how context modulates natural sound selectivity by showing that previous acoustic sequences have disparate effects on neuronal discriminability. The presence of leading context turns bad natural-sound discriminators into good ones and has the opposite effect on good discriminators. Moreover, such modulation appears to be relative to neuronal type (i.e. putative pyramidal vs. interneuron see Fig. 3b). These findings may have bearings on our interpretation on how the auditory cortex operates in natural conditions. We propose that non-selective neurons are important for coding violations to regularities in the sound stream, i.e. transitions from echolocation to communication (or vice versa). Meanwhile, neurons that appear to be selective to specific natural sound categories when there is no context would actually be worse detectors of acoustic transitions, but this population of neurons is probably key for sound discrimination in silent environments. Future studies could test whether this occurs in other mammalian species with less specialized hearing compared to bats.

### Mechanisms underlying context-specific response modulation

We show that in context-sensitive neurons, the presence of a leading context always reduced responses to forthcoming sounds independently of the neuronal tuning. We propose that there is a common mechanism that underlies such context effect in cortical neurons of non-tonotopic areas. Reduction of the responsiveness after stimulation has usually been attributed to synaptic (GABAergic) inhibition as well as long-lasting mechanisms, such as synaptic depression (Wehr & Zador, 2005; Asari & Zador, 2009). Such mechanisms have been described to be activity-dependent, operating at the level of the neuron’s output (Calford & Semple, 1995; Brosch & Schreiner, 1997; Wehr & Zador, 2005) or input (Ulanovsky *et al*., 2003).

Our model predicts that context modulation in neuronal responses to natural sounds results from two different mechanisms: adaptation dependent on the postsynaptic activity and adaptation dependent on the presynaptic activity. The model assumes that the processing of natural sounds is segregated in different frequency channels that converge into a cortical neuron that exhibits context-sensitivity. Thus, hearing the context unbalances the input-dependent adaptation and allows context-specific effects. Similar mechanisms, such as synaptic depression, have been described to explain specific suppression of repetitive artificial sounds (Chung *et al*., 2002; Eytan *et al*., 2003; Wehr & Zador, 2005; Rothman *et al*., 2009). However, the suppression of *unexpected* sounds cannot be explained by frequency specific adaptation alone. We predict that postsynaptic-dependent adaptation, which attenuates every input to the neurons, is needed to explain the totality of our results. Probably potassium currents are involved in this type of mechanism (Abolafia *et al*., 2011). Previous modelling studies of ‘adaptation channels’ have explained history-dependent effects in the neuronal response to artificial sounds but using the oddball paradigm (Puccini *et al*., 2006; Taaseh *et al*., 2011; May *et al*., 2015).

We should point out that in our model, the synaptic depression is assumed to occur in excitatory synapses arriving to cortical cells. However, inhibitory synapses could also reproduce our predictions. Indeed, the role of GABAergic neurotransmission has been demonstrated in similar context-dependent situations (Perez-Gonzalez *et al*., 2012). It remains a challenge for future work to discover the biological correlate(s) of the current model synapses. Whether the synaptic depression depends on the activity of thalamic or cortical neurons remains unsolved. With our data we also cannot establish whether the effects observed originate at the cortical level, or whether are simply inherited by cortical neurons. What we do propose based on our model is that, regardless of the input origin, the selectivity to natural sounds in the presynaptic sites is tuned to the spectral components of natural sounds, more than to sound categories (response either to echolocation or communication signals). Our results showed that preference to natural sounds, but not exclusivity, is sufficient to reproduce and better fit the observed context-specific modulatory effects. The latter is in accordance with several reports in lower areas of the auditory pathway and in cortical areas, where the neurons exhibit high selectivity to natural sounds (Klug *et al*., 2002; Mayko *et al*., 2012; Salles *et al*., 2020).

In summary, our results indicate that acoustic context has strong effects on cortical neuronal discriminability of natural sounds. These effects turn neurons from non-tonotopic areas into good discriminators of natural sound transitions, and they can be explained based on multiple input channels that display adaptation and that are differentially tuned to the spectral components of natural vocalizations.

## Methods

### Animal preparation

Six bats (2 females, species *Carollia perspicillata*) were used in this study. They were taken from a breeding colony in the Institute for Cell Biology and Neuroscience at the Goethe University in Frankfurt am Main, Germany. All experiments were conducted in accordance with the Declaration of Helsinki and local regulations in the state of Hessen (Experimental permit #FU1126, Regierungspräsidium Darmstadt).

The bats were anesthetized with a mixture of ketamine (100 mg/ml Ketavet; Pfizer) and xylazine hydrochloride (23.32 mg/ml Rompun; Bayer). Under deep anaesthesia, the dorsal surface of the skull was exposed. The underlying muscles were retracted with an incision along the midline. A custom-made metal rod was glued to the skull using dental cement to fix the head during electrophysiological recordings. After the surgery, the animals recovered for 2 days before participating in the experiments.

On the first day of recordings, a craniotomy was performed using a scalpel blade on the left side of the cortex in the position corresponding to the auditory region. Particularly, the caudoventral region of the auditory cortex was exposed, spanning primary and secondary auditory cortices (AI and AII, respectively), the dorsoposterior field (DP) and high-frequency (HF) fields. The location of these areas was made by following patterns of blood vessels and the position of the pseudocentral sulcus (Esser & Eiermann, 1999; Hagemann *et al*., 2010).

### Neuronal recordings

In all six bats, recordings were performed over a maximum of 14 days. Experiments were conducted in awake animals. Bats were head-fixed and positioned in a custom-made holder over a warming pad whose temperature was set to 27°C. Local anaesthesia (Ropivacaine 1%, AstraZeneca GmbH) was administered topically over the skull before each session. Each recording session lasted a maximum of 4h.

All experiments were performed in an electrically isolated & sound-proofed chamber. For neural recordings, carbon-tip electrodes (impedance ranged from 0.4 to 1.2 MΩ) were attached to an electrode holder connecting the electrode with a preamplifier to a DAGAN four channel amplifier (Dagan EX4-400 Quad Differential Amplifier, gain = 50, filter low cut = 0.03 Hz, high cut = 3kHz). A/D conversion was achieved using a sound card (RME ADI-2 Pro, SR = 192 kHz). Electrodes were driven into the cortex with the aid of a Piezo manipulator (PM 10/1; Science Products GmbH). Single unit auditory responses were located at depths of 308 +-79 μm, mean +-SD, using a broadband search stimulus (downward frequency modulated communication sound of the same bat species) that triggered activity in both low and high frequency tuned neurons. A similar paradigm has been used in previous studies to locate neuronal responses (Martin *et al*., 2017).

### Acoustic stimulation

We used natural sounds to trigger neural activity during the recordings. The natural sounds were obtained from the same species in previous studies from our lab (Beetz *et al*., 2016; Hechavarria *et al*., 2016). Acoustic signals were generated with an RME ADI.2 Pro Sound card and amplified by a custom-made amplifier. Sounds were then produced by a calibrated speaker (NeoCD 1.0 Ribbon Tweeter; Fountek Electronics, China), which was placed 15 cm in front of the bat’s right ear. The speaker’s calibration curve was calculated with a microphone (model 4135; Brüel & Kjaer).

Once an auditory neuron was located, we determined the iso-level frequency tuning of the unit, with 20-ms pure tones (0.5 ms rise/fall time) presented randomly in the range of frequencies from 10 to 90 kHz (5 kHz steps, 20 trials) at a fixed sound pressure level of 60 dB SPL. This was done only in a subset of the neurons recorded (55 out of a total of 74). Our previous studies indicate that 60 dB SPL represents a good compromise since it is strong enough to drive activity in most auditory cortex (AC) neurons and allows to differentiate between single-peaked and double-peaked tuning curves typical of the AC of this bat (Lopez-Jury *et al*., 2020). The repetition rate for stimulus presentation was 2 Hz.

We studied context-dependent auditory responses using two types of acoustic contexts: sequences of echolocation calls and sequences of distress calls. We refer to these two sequences of natural vocalization as “context” because they preceded the presentation of probe sounds that were either a single echolocation or a single distress syllable. The properties of the contexts used for stimulation are depicted in Fig. 1a-b. In a nutshell, the echolocation sequence was composed of high-frequency vocalizations and their echoes (carrier frequencies > 50 kHz) repeated at intervals ~ 40 ms (Fig. 1a-b left panels). The echolocation sequence was recorded from a bat swung in a pendulum following procedures described elsewhere (Beetz *et al*., 2016). The distress sequence, on the other hand, was composed of individual syllables with peak frequencies ~ 23 kHz (Fig. 1a-b right panels). Distress syllables occurred in groups (so-called bouts) repeated at intervals ~ 60 ms and within the bouts, syllables occurred at rates ~ 15 ms (Hechavarria *et al*., 2016). We chose to study these two acoustic contexts because they rely on fundamentally different acoustic parameters and are linked to distinct behaviours, i.e. navigation (echolocation) and calling under duress (distress).

Probe sounds (sounds that followed the context) were single echolocation and distress syllables, each obtained from the respective context sequences (Fig. 1c). We tested two temporal separations (gaps) between the context offset and probe onset: 60 and 416 ms. Therefore, a total of 8 context-stimuli (2 contexts x 2 probes x 2 gaps) were randomly presented and repeated 20 times to awake bats during electrophysiological recordings: navigation context followed by navigation probe; navigation context followed by distress probe, distress context followed by navigation probe and distress context followed by distress probe. In addition, we presented each probe after 3.5 s of silence (no context).

### Data analysis

#### -Spike clustering

All the recording analyses, including spike sorting, were made using custom-written Matlab scripts (R2018b; MathWorks). The raw signal was filtered between 300 Hz and 3 kHz using a bandpass Butterworth filter (3^rd^ order). To extract spikes from the filtered signal, we detected negative peaks that were at least three standard deviations above the recording baseline; times spanning 1 ms before the peak and 2 ms after were considered as one spike. The spike waveforms were sorted using an automatic clustering algorithm, “KlustaKwik,” that uses results from PCA analyses to create spike clusters (Harris *et al*., 2000). For each recording, we considered only the spike cluster with the highest number of spikes.

#### -Neuronal classification

A total of 74 units were considered as responsive to at least one of the probe sounds tested. A unit was considered as responsive if the number of spikes fired in response to the sound in question was above the 95% confidence level calculated for spontaneous firing for the same unit (calculated along 200 ms before the start of each trial). Evoked firing had to surpass this threshold for at least 8 ms after probe onset for a unit to be considered as responsive. To test for natural sound-preferences in each unit (i.e. whether units preferred echolocation vs. distress sounds or vice versa), we used the responses to the probe sounds when they were preceded by silence (no context). The spike counts during 50 ms after each probe onset were compared using a non-parametric effect size metric: Cliff’s delta. Cliff’s delta quantifies effect size by calculating how often values in one distribution are larger than values in a second distribution. Distribution here refers to the number of spikes fired in presentations of navigation and distress probe trials. The statistic gives values from −1 to 1, with identical groups rendering values of zero. Following previous studies (Romano *et al*., 2006), if the effect size between the two distributions was negligible or small (abs(Cliff’s delta)<=0.3), the unit was classified as ‘equally responsive’ to both probes. On the other hand, if abs(Cliff’s delta)>0.3, the unit was classified either as ‘preference to echolocation’ or ‘preference to distress’. The preference was assigned to the probe that evoked the highest number of spikes across trials. If a unit was responsive to only one of the probes, it was classified as having preference for ‘only echolocation’ or ‘only distress’. We checked the classification criterion by using non-parametric Wilcoxon rank-sum tests. Rank-sum test failed to reject the null hypothesis when comparing spike counts of the responses to both probes in all the units classified as ‘equally responsive’ based on the Cliff’s delta metric. In contrast, in all sixteen units considered as ‘preference to distress’ the null hypothesis was rejected, i.e. they showed significant differences in spike counts across trials (p-value < 0.05, Wilcoxon rank-sum test).

The iso-level tuning curves were obtained from the average of spike counts across trials in response to pure tones at *N* frequencies, rescaled by min-max normalization:

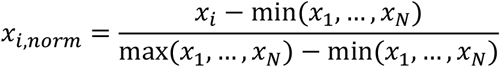

Units were classified as low-frequency tuned if the normalized response was higher than or equal to 0.6 at any frequency lower than 50 kHz and lower than 0.6 for all frequencies higher than or equal to 50 kHz. High-frequency tuned neurons were those in which *x_i,norm_* was above 0.6 for any frequency higher than or equal to 50 kHz and lower than 0.6 for all of the frequencies lower than 50 kHz. Multi-peaked neurons were those in which *x_i,norm_* exceeded 0.6 in both frequency bands (< 50 kHz and >= 50 kHz). A similar classification has been used in previous studies (Lopez-Jury *et al*., 2020).

#### -Quantifying the effects of leading acoustic context

To test for effects of leading acoustic context on probe-triggered responses, we tested (using a non-parametric Wilcoxon rank-sum test) whether the response to the probe, in terms of number of spikes across trials, occurring after the context was significantly different from the responses to the same sound presented without any context. To quantify the magnitude of the effect, we used the following equation:

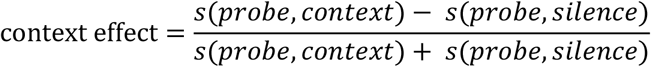

where *s*(*probe, context*) corresponds to the number of spikes during 50 ms from the onset of the corresponding probe following the corresponding context and *s*(*probe, silence*), following silence. Effects of the same context on different probe-triggered responses were statistically compared per unit using paired statistics (Wilcoxon signed rank test). In addition, we compared the calculated effect size (Cliff’s delta test) between probes after silence with the same measurement after context, using also paired statistics (Wilcoxon signed rank test). According to our convention, negative Cliff’s delta values indicate higher responses to the distress probe than to the echolocation probe. Positive Cliff’s delta values indicate the opposite. For space reasons, the results presented in this paper focus only to data obtained using a temporal gap of 60 ms between context offset and probe onset. Data obtained with 416-ms gaps rendered similar results (Supplementary Fig. 2).

#### Modelling possible synaptic origins of contextual neural response modulation

We modelled a broadly-tuned cortical neuron that reproduces the behaviour of ‘equally’ responsive neurons observed in our experiments, using an integrate-and-fire neuron model. Our model (described in detail below) receives input from two narrow frequency channels and includes synaptic adaptation (i.e. activity-dependent depression) in the cortical input synapses.

Frequency specific adaptation in multiple synaptic channels has been proposed to mediate stimulus-specific adaptation in auditory cortex, including during speech processing (Taaseh *et al*., 2011; May & Tiitinen, 2013). All simulations were performed using the Python package Brian 2 version 2.3 (Stimberg *et al*., 2019).

In the model proposed, the neuron’s membrane potential (*V_m_*) evolves according to the following differential equation:

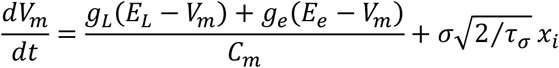

where *g_L_* is the leak conductance, *E_L_* is the resting potential, *C_m_* is the membrane capacitance, *E_e_* is the excitatory reversal potential and *g_e_* is the synaptic conductance. The second term of the equation corresponds to stochastic current added to the neuron model. *x_i_* is a Gaussian random variable specified by an Ornstein-Uhlenbeck process, with variance *σ* and correlation time *τ_σ_*.

The neuron fires when *V_m_* reaches the membrane potential threshold *ω_th_* and the variable *V_m_* is reset to the fixed value *V_r_*. To implement a reduction of the firing in response to constant stimulation, we added an adaptive threshold to the model that decreases the probability of firing with postsynaptic activity. The firing threshold *ω_th_* starts at *V_th_* and after the occurrence of a spike, it is increased by the amount Δ_*th*_ and it returns back to *V_th_* with a time constant *τ_th_*, that is,

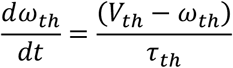

The model includes conductance-based excitatory synapses, every synapse creates a fluctuation of conductance *g_e_* that change in time with time constant *τ_e_*, as follow,

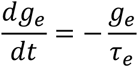

When an action potential arrives at a synapse, the excitatory conductance *g_e_* increases in the postsynaptic neuron, according to

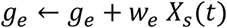

where *W_e_* is the maximum synaptic weight and is modulated by a time-dependent adaptation term *X_s_*(*t*). The effective synaptic weight between the inputs and the cortical neuron was the product *w_e_ X_s_*(*t*) and depended on the presynaptic activity. The model assumes that *X_s_* starts at one and a presynaptic spike arrival at synapse induces a decrement of *X_s_* value by the amount Δ_*s*_, which recovers in time by:

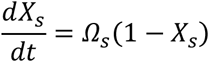

The neuron model has two excitatory synapses that receive inputs, which differ by their preference for natural sounds stimulation. The input of the synapse *j* corresponds to a spike train that follows an inhomogeneous Poisson process with rate *λ_j_*:

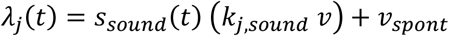

where *s_sound_*(*t*) corresponds to the envelope in time of the respective natural sound, *k_j,sound_* is proportional to the response of input *j*to the respective sound, *υ* is the average firing rate of the inputs and *υ_spont_* is the spontaneous firing rate of the inputs.

#### Model parameters and data fitting

All the parameters used in the model are indicated in Table 1. The parameters associated to intrinsic properties of the neuron model were set to qualitatively mimic the extracellular activity of cortical neurons. The parameters that determine the dynamic of the adaptive threshold (Δ_*th*_ and *τ_th_*) were set to produce a reduction of 40-50% in the response to the probes after any context, as observed empirically.

**Table 1.** Parameters used in the neuron model.

| Parameters | Values |
| --- | --- |
| *Neuron model* |  |
| $C_m$, membrane capacitance | 100 pF |
| $E_L$, leak reversal potential | -55 mV |
| $g_L$, leak conductance | 5 nS |
| $V_{th}$, firing threshold potential | -50 mV |
| $V_r$, reset potential | -55 mV |
| $\sigma$, sigma noise | 2 mV |
| $\tau_\sigma$, time constant noise | 10 ms |
| *Neuronal adaptation* |  |
| $\Delta_{th}$, adaptive threshold increment | 0.005 mV |
| $\tau_{th}$, adaptive threshold time constant | 5 s |
| *Excitatory synapses* |  |
| $E_e$, excitatory reversal potential | 0 mV |
| $\tau_e$, excitatory time constant | 10 ms |
| $w_e$, synaptic weight | 6 nS |
| *Input* |  |
| $v$, average input rate | 2 / ms |
| $v_{spont}$, spontaneous input rate | 1 / s |
| $k_{i,com}$, response input $i$ to communication | 0.7 |
| $k_{j,com}$, response input $j$ to communication | 0.1 |
| $k_{i,nav}$, response input $i$ to navigation | 0 |
| $k_{j,nav}$, response input $j$ to navigation | 1.5 |
| *Synaptic adaptation* |  |
| $\Omega_{i,s}$, adaptation recovery rate input $i$ | 1.9 / s |
| $\Omega_{j,s}$, adaptation recovery rate input $j$ | 1.4 / s |
| $\Delta_{i,s}$, synaptic decrement input $i$ | 0.0125 |
| $\Delta_{j,s}$, synaptic decrement input $j$ | 0.025 |

**Table 2.** Parameters that were modified from the fitted neuron model.

| Models varying degree of input selectivity |  |  |
| --- | --- | --- |
| *No* |  |  |
| $k_{i,com}$ | response input $i$ to communication | 0.4 |
| $k_{j,com}$ | response input $j$ to communication | 0.4 |
| $k_{i,nav}$ | response input $i$ to navigation | 0.8 |
| $k_{j,nav}$ | response input $j$ to navigation | 0.8 |
| *High* |  |  |
| $k_{i,com}$ | response input $i$ to communication | 0.8 |
| $k_{j,com}$ | response input $j$ to communication | 0 |
| $k_{i,nav}$ | response input $i$ to navigation | 0 |
| $k_{j,nav}$ | response input $j$ to navigation | 1.5 |
| *Low* |  |  |
| $k_{i,com}$ | response input $i$ to communication | 0.7 |
| $k_{j,com}$ | response input $j$ to communication | 0.1 |
| $k_{i,nav}$ | response input $i$ to navigation | 0 |
| $k_{j,nav}$ | response input $j$ to navigation | 1.5 |
| Models varying adaptation |  |  |
| *None* |  |  |
| $\Delta_{th}$ | adaptive threshold increment | 0 mV |
| $\Delta_{i,s}$ | synaptic decrement input $i$ | 0.0 |
| $\Delta_{j,s}$ | synaptic decrement input $j$ | 0.0 |
| *Post* |  |  |
| $\Delta_{th}$ | adaptive threshold increment | 0.005 mV |
| $\Delta_{i,s}$ | synaptic decrement input $i$ | 0.0 |
| $\Delta_{j,s}$ | synaptic decrement input $j$ | 0.0 |
| *Pre* |  |  |
| $\Delta_{th}$ | adaptive threshold increment | 0 mV |
| $\Delta_{i,s}$ | synaptic decrement input $i$ | 0.0125 |
| $\Delta_{j,s}$ | synaptic decrement input $j$ | 0.025 |
| *Both* |  |  |
| $\Delta_{th}$ | adaptive threshold increment | 0.005 mV |
| $\Delta_{i,s}$ | synaptic decrement input $i$ | 0.0125 |
| $\Delta_{j,s}$ | synaptic decrement input $j$ | 0.025 |

In our model the reversal potential *E_e_* of excitatory synapses was set to a standard value used for modelling AMPA synapses (Clopath *et al*., 2010). The time constant *τ_e_* was adjusted to obtain spiking responses to the probes qualitatively similar in duration with the observed experimental data.

The maximum synaptic weight *w_e_* of each synapse was set in order to fit experimental data. We ran 81 simulations systematically changing the maximum synaptic weight of each synapse independently from 1 to 9 nS, in steps of 1 nS. For each simulation we compared statistically (using Wilcoxon rank-sum test) 50 neuron models with the 45 units classified as ‘equally’ responsive, regarding Cliff’s delta values between probe-responses (Supplementary Fig. 8a1). In addition, the chosen parameters were constraint to fit approximately the number of spikes evoked by each probe observed empirically (Supplementary Fig. 8a2).

We assumed that the spiking of the inputs was tuned to the spectral content of natural sounds: high-frequency tuned (>=45 kHz) and low-frequency tuned (<45 kHz). Considering the spectral distribution of the vocalizations used as context (Fig. 1b), high-frequency tuned input was set to be responsive to biosonar signals as well as to distress syllables, and low-frequency tuned input to only distress. The input selectivity to natural sounds is given by the parameter *k_j,sound_*. We tested several ratios of selectivity to distress syllables between the inputs (*k_low,distress_/k_high,distress_*), ranging from equally responsive to low-frequency input exclusively responsive. The ratios were tested for several synaptic decrement values (Supplementary Fig. 8b) since the behaviour of the model is highly dependent on this parameter. Ratios close to 1 were unable to reproduce our data after context, especially after the communication sequence. We chose a ratio of 0.7/0.1 because it is similar to the proportion of low frequencies components relative to high frequencies in distress calls. The average rate of the inputs was calibrated to generate spike trains with one or two spikes during the probe stimulation (~1.5 ms of duration).

Our model includes short-term synaptic adaptation (i.e. activity-dependent depression) that depends essentially on two parameters: the synaptic decrement Δ_*s*_ and the adaptation recovery rate *Ω_s_*. The interplay between these two parameters was systematically tested by changing them on each synapse. Results in terms of Cliff’s delta values obtained between the two probes are shown in Supplementary Fig. 5a-b. The chosen values do not show significant differences with experimental data after both contexts. To check whether the chosen parameters fit the data independently of the synaptic weight of the synapses, we ran several simulations changing the maximum synaptic weight of each synapse. No matter which value was tested, the discriminability index was always lower after navigation context and higher after communication context in comparison with the values obtained after silence (Supplementary Fig. 5c).

Finally, we modified our model in order to reproduce experimental data obtained in units classified as ‘preference to distress’. We ran several simulations changing the maximum synaptic weight *w_e_* of each synapse (Supplementary Fig. 7a). A higher synaptic weight in the low-frequency synapse and lower weight in the high-frequency synapse turned our neuron model into a ‘preference to distress’ unit. Without changing any other parameter, we observed that the model reproduces the behaviour of these neurons after context (Supplementary Fig. 7b).

## Author contributions

LLJ, JCH, FGR, and EGP designed research; LLJ performed experiments and analyzed data; LLJ wrote the original draft; JCH, FGR and EGP reviewed and edited the draft.

