## supplementary figures for "Acoustic context modulates natural sound discrimination in auditory cortex through frequency specific adaptation"

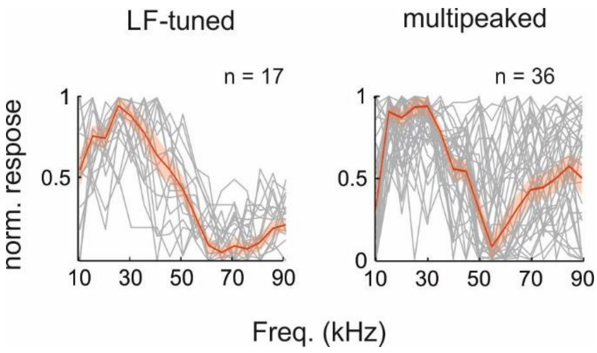


**Supplementary Figure 1.** Iso-level tuning curves from units classified as low-frequency tuned (left) and from units classified as 'multipeaked' (right). The red line indicates the average of the corresponding grey curves and the shade area the respective standard error. Note that the peak on lower frequencies of multipeaked tuning curves is more consistent across units than the first peak.

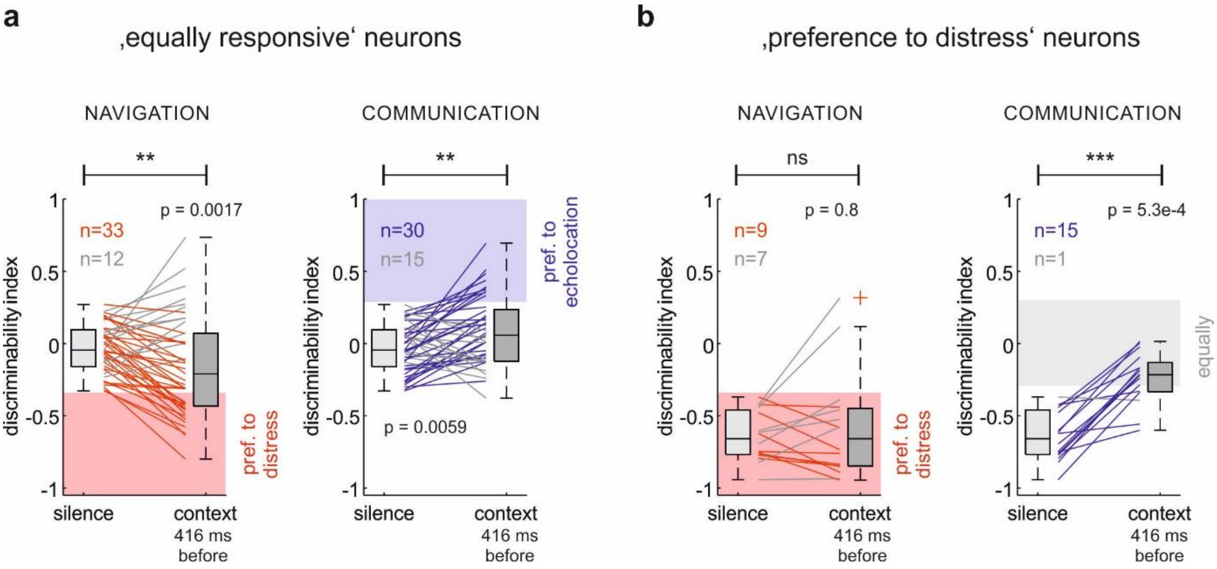


**Supplementary Figure 2. Similar results from data obtained using a temporal gap of 416 ms between context offset and probe onset.**

**a** Cliff's delta of 'equally responsive' units comparing echolocation and distress responses after silence and after acoustic context. Note that both contexts increase neuronal discriminability. **b** Cliff's delta of 'preference for distress' units comparing responses to echolocation and distress syllables after silence and after acoustic context. Note that, in this case, communication context decreases neuronal discriminability. In all the plots, the significance level is indicated above; *p*-values were obtained using the paired test Wilcoxon signed rank.

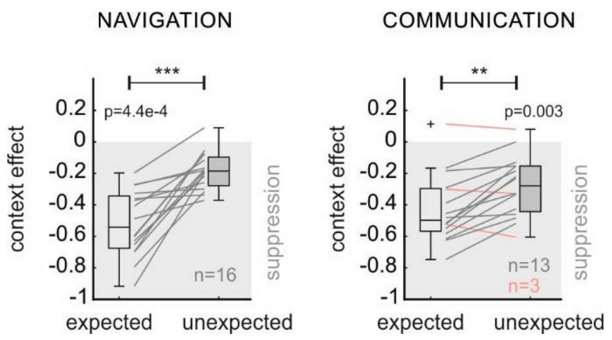


**Supplementary Figure 3.** Context effect quantification of ‘preference for distress’ units for *expected* and *unexpected* probes. Lines join values from the same neuron. Grey lines correspond to units with a higher suppression on *expected* sounds. Otherwise, the lines are pink. Significance level is indicated above the plots; *p*-values were obtained using the paired test Wilcoxon signed rank.

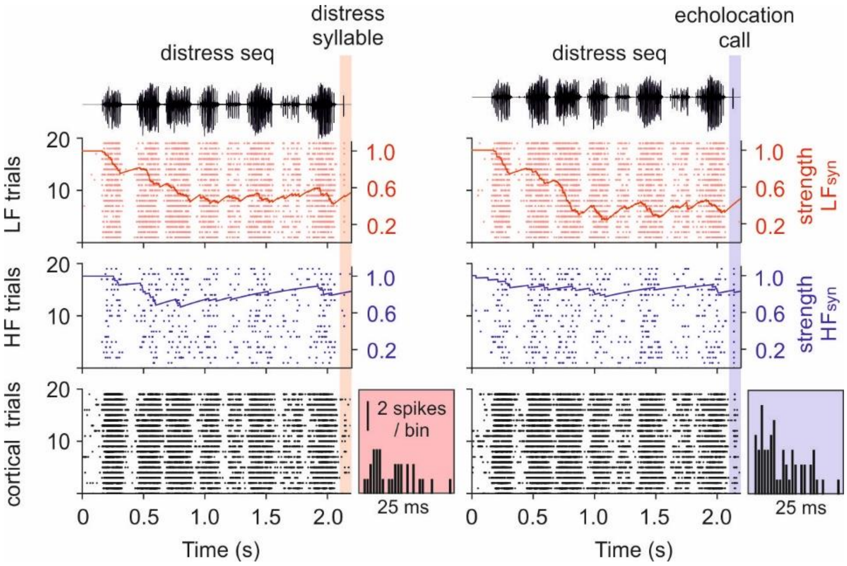


**Supplementary Figure 4.** Example of 20 simulations of neuronal responses to a distress sequence followed by a distress syllable (left) and by an echolocation call (right). Raster plots indicate the spiking of the inputs (top and middle) and of the cortical neuron model (bottom). Together with the rasters, it is also depicted the time evolution of the synaptic strength associated to the spiking of the first trial of each input. The insets show the PSTH obtained from the neuron model during 25 ms from the onset of the probe for each stimulation.

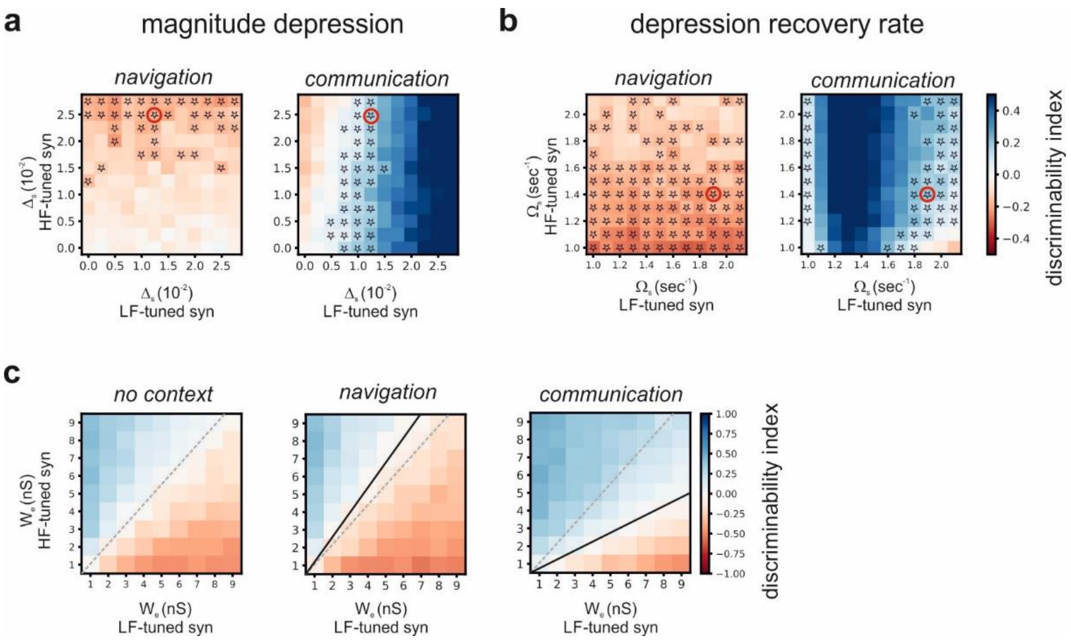


**Supplementary Figure 5. Fitting the synaptic adaptation to experimental data.**

**a** Fitting the magnitude of the synaptic depression of the model to experimental data. Average discriminability index of 50 units obtained from 144 simulations systematically changing the synaptic decrement ($\Delta_S$) of LF tuned and HF tuned synapses (from 0 to 0.0275, in steps of 0.0025); after navigation context (left) and after communication context (right). **b** Same than in A but changing the recovery rate of the synaptic depression ($\Omega_S$), from 1.0 /s to 2.1 /s, in steps of 0.1. Stars indicate non-significant differences between the model and our experimental data using the non-parametric Wilcoxon rank sum test (*p*-value>0.05). Red circles indicate the combination of parameters chosen for the model reproduce our experimental data. **c** Average discriminability index of 50 units (modelled with the chosen parameters indicated in A-B) obtained from 81 simulations systematically changing the maximum synaptic weight ($w_e$) of LF tuned and HF tuned synapses (from 1 nS to 9 nS, in steps of 1), after the context indicated above of each plot. Dashed grey lines indicate discriminability index equal to zero when no context preceded the probes. The black lines indicate also null discriminability indices but after context and illustrate the shift of the indexes towards left, after a navigation context and towards right, after a communication context.

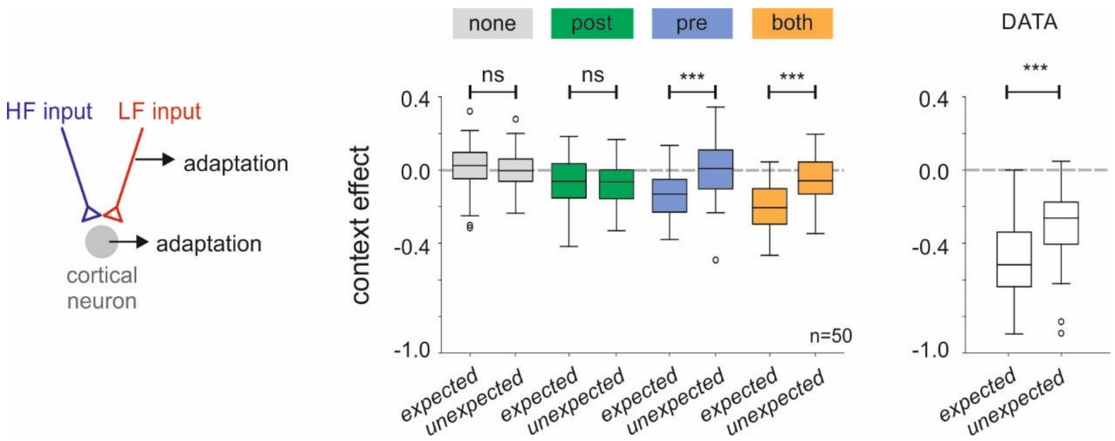


**Supplementary Figure 6.** Navigation context effect calculated for *expected* and *unexpected* probes using models with different forms of adaptation. ‘None’: model without any type of adaptation. ‘Post’: model with neuronal adaptation. ‘Pre’: model with synaptic adaptation. ‘Both’: model with neuronal and synaptic adaptation. Right boxplots show the respective context effect obtained from experimental data. In all the comparisons, the significance level is indicated above; *p*-values were obtained using non-paired Wilcoxon rank sum test for the simulations, and the paired Wilcoxon signed rank for the real data. The parameters used on each model are indicated in Table 2.

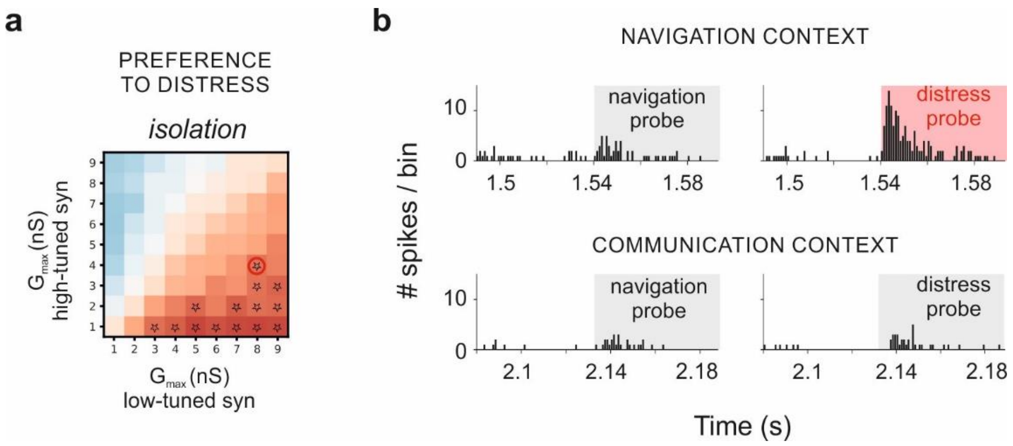


**Supplementary Figure 7. ‘Preference to distress’ neuron model.**

**a** Average of discriminability index after no context of 50 units obtained from 81 simulations systematically changing the maximum synaptic weight ($W_e$) of of low-frequency (LF) tuned and high-frequency (HF) tuned synapses (from 1 nS to 9 nS, in steps of 1). Stars indicate non significative differences between the model and our experimental data (specifically for ‘preference for distress’ units, n=16) using the non-parametric Wilcoxon rank sum test (*p*-value>0.05). Red circle indicates the combination of parameters chosen to the model reproduce the data obtained from this neuron category. **b** Simulated responses to each probe after a sequence of echolocation (top) and after a distress call (bottom).

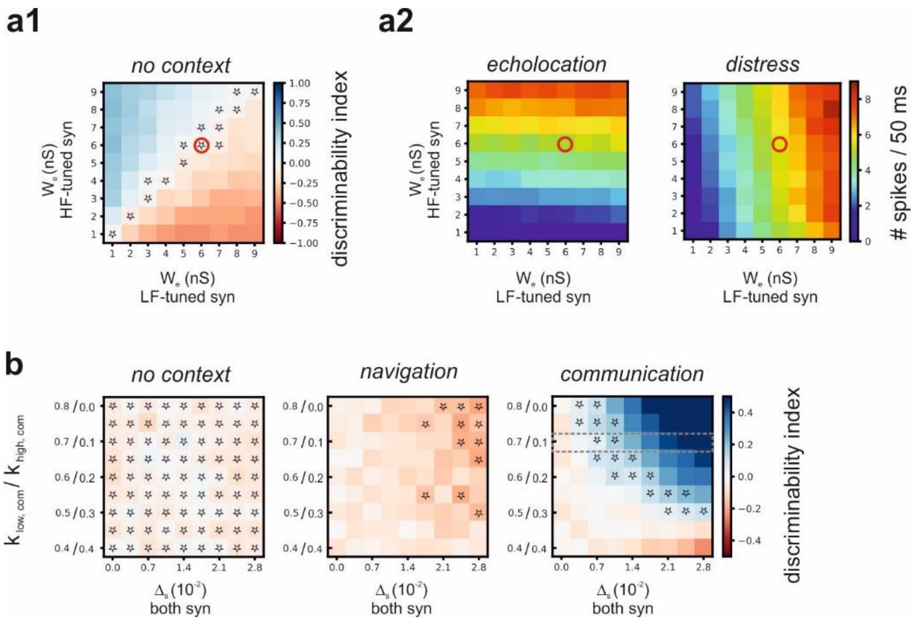


**Supplementary Figure 8. Fitting the synaptic weight of synapses and the degree of input selectivity to experimental data.**

**a1** Average discriminability index after no context of 50 units obtained from 81 simulations systematically changing the maximum synaptic weight ($w_e$) of low-frequency (LF) tuned and high-frequency (HF) tuned synapses (from 1 nS to 9 nS, in steps of 1). **a2** Average number of spikes during 50 ms after the onset of the probe (echolocation call at left and distress syllable at the right) obtained from the same 81 simulations showed in A1. Red circles indicate the combination of parameters chosen to the model reproduce our data. **b** Average discriminability index of 50 units obtained from 81 simulations systematically changing the degree of selectivity of LF tuned and HF tuned input for distress syllables. The ratio ranges from 0.4/0.4 to 0.8/0.0, increasing the numerator in steps of 0.05. For each ratio we tested 9 magnitudes of synaptic depression, ranging from 0 to 0.028, in steps of 0.0035. Each plot was obtained with different contexts, indicated on top of each. Dashed grey rectangle indicate the ratio chosen to the model reproduce our data. Stars indicate non-significant differences between the model and our experimental data using the non-parametric Wilcoxon rank sum test ($p$-value>0.05).
